# Production of membrane-embedded Bcl-2 proteins - Use of cell-free synthesis in continuous exchange for co-translational insertion of Bcl-2 proteins in lipid bilayer nanodiscs

**DOI:** 10.64898/2026.08.15.745005

**Authors:** Joann Kervadec, Akandé Rouchidane Eyitayo, Cécile Gonzalez, Thomas Maurice, Karine Bernardeau, Stephen Manon, Muriel Priault

## Abstract

The BCL-2 family proteins are key regulators of apoptosis, functionally divided in pro- and anti-apoptotic proteins, with a third group acting as regulators. Their ability to partition between the cytosol and intra-cellular membranes (essentially the mitochondrial outer membrane) is a primary regulator of their functions. A second contributor is their ability to form homotypic complexes (pro-pro or anti-anti) or heterotypic complexes (pro-anti). If the structures of monomeric cytosolic members have largely been characterized, the functional and structural study of membrane-embedded proteins remains incomplete. Unlocking this knowledge is expected to enable evaluating new therapeutic strategies to either activate pro-apoptotic members, or inactivate anti-apoptotic ones. Lipid bilayer nanodiscs and improved cell-free protein synthesis have provided the technical breakthrough to achieve the description at the atomic level of conformations and higher order assemblies of these proteins in their membrane-associated states. Here we describe detailed and straightforward protocols for generating nanodisc-inserted members of the Bcl-2 family, through the example of anti-apoptotic Bcl-xL, and pro-apoptotic Bax and Bak. Full-length, untagged proteins are expressed from bacterial extracts in the presence of pre-assembled nanodiscs to allow co/post-translational insertion in lipid bilayer, followed by affinity chromatography purification. A more detailed characterization is presented for Bak, to exemplify structural and mechanistic studies enabled by these methods.

**Graphical abstract:** 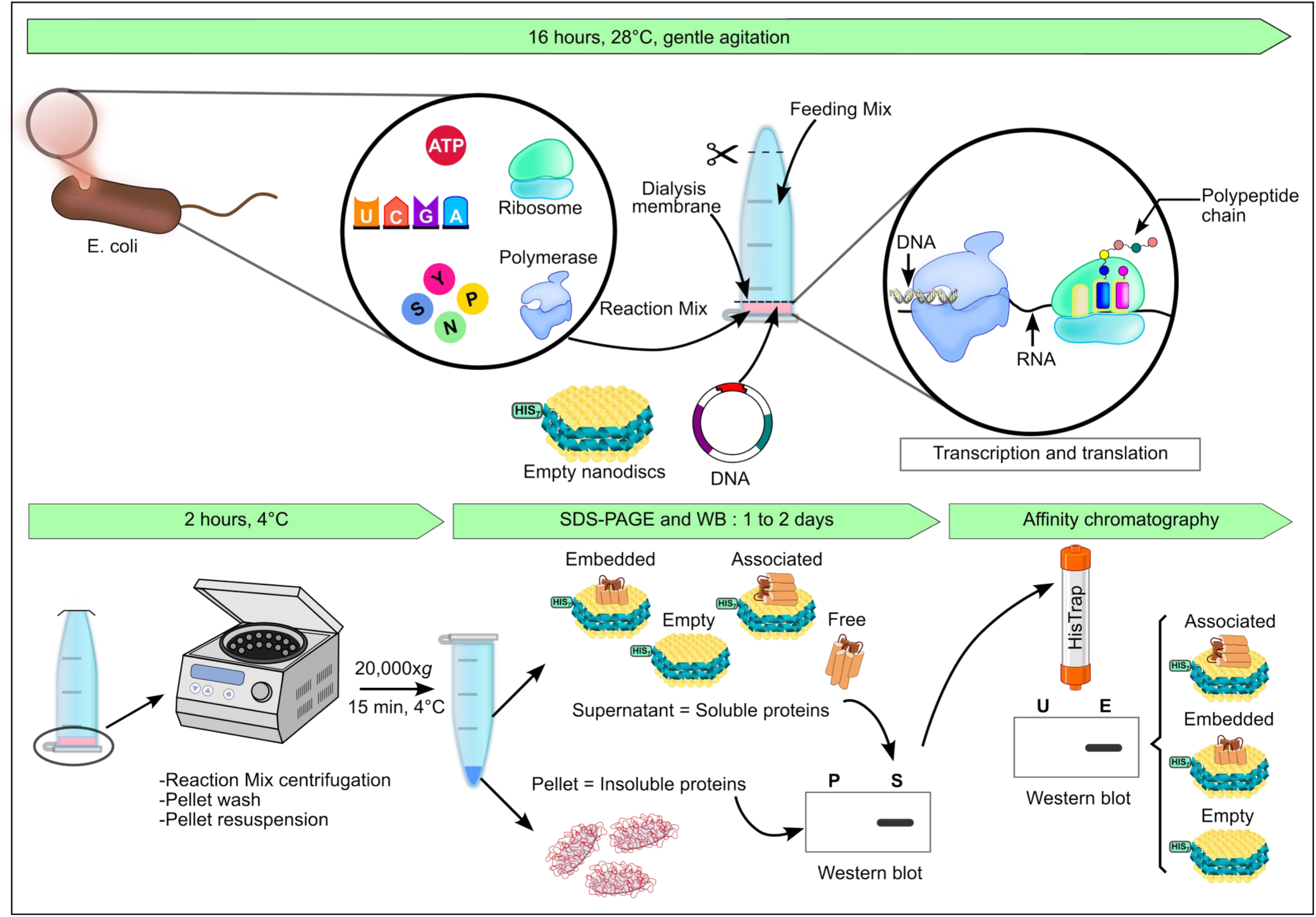

## 1. Introduction

Bcl-2 family proteins are involved in the control of the mitochondrial pathway leading to apoptosis, a highly regulated form of programmed cell death in metazoans. These small globular proteins (21 to 38 kDa) share <u>B</u>cl-2 <u>H</u>omology motives numbered from BH1 to BH4, and are subdivided in two functional subgroups. A first subgroup comprises multi-motif anti-apoptotic members, such as Bcl-xL and Bcl-2. They inhibit a second subgroup of multi-motif pro-apoptotic members, such as Bax and Bak, through well characterized canonical interactions; pro-apoptotic members also include BH3-only proteins, which either directly activate multi-motif pro-apoptotic proteins, or release inhibition by anti-apoptotic proteins (for review, see (Czabotar & Garcia-Saez, 2023)).

Stepwise conformational changes allow multi-domain Bax and Bak to activate and transition from the cytosol to mitochondrial outer membranes (MOM), where they oligomerize and create pores that release apoptogenic proteins toward the cytosol to amplify apoptosis signaling. Inhibitory interactions with Bcl-xL occur in the cytosol and prevent MOM permeabilization, but it is still unclear whether interactions persist at the mitochondrial membrane, and if so, what supramolecular arrangements are formed between pro-and anti-apoptotic members. Bcl-2 proteins are important drug targets in cancer therapy, as anti-apoptotic proteins are often over-expressed in various cancer types. Fundamental knowledge was successfully transferred to the clinic when synthetic BH3-mimetic molecules proved efficient to disrupt the inhibitory interactions between anti-apoptotic BH1-BH2-BH3 hydrophobic groove and the BH3 motif of pro-apoptotic partners (Lessene et al., 2013). However, the converse strategy to design drugs that activate pro-apoptotic proteins is still hindered by insufficient information on the active structures and supramolecular arrangements of membrane-embedded Bcl-2 proteins.

Recently, two technical improvements have significantly facilitated protein study *in vitro* : the refinement of cell-free protein production on the one hand, and the use of artificial nano-membranes called nanodiscs on the other hand. Cell-free protein synthesis (CFPS) consists in using the bacterial transcription-translation machinery to produce *in tubo* a given protein from its cDNA cloned in a plasmid. Former “Rapid Translation System” has enabled recombinant protein production for over 20 years, but the recent refinement was the yield increase to mg of proteins, thanks to the continuous exchange of substrates and reaction products through a dialysis membrane (Schwarz et al., 2007), followed by incremental optimizations (Dopp et al., 2019). Nanodiscs (NDs) are non-covalent discoidal bilayers of phospholipid stabilized by apolipoprotein-derived helical membrane scaffold proteins (MSP) (Civjan et al., 2003). NDs are highly tractable tools allowing to adjust lipid composition (polar heads, acyl chain unsaturation and length) as well as their diameter, thanks to a versatile library of genetically engineered MSPs (Denisov et al., 2004). Moreover, MSP proteins may harbor affinity tags to enable purification. The use of NDs renders membrane proteins soluble in aqueous solutions without detergents, and the native-like bilayer environment ensures protein stability and maintains activity. The combination of CFPS and NDs has already allowed co-translational insertion of Bcl-xL and Bax by our team (Rouchidane Eyitayo, Boudier-Lemosquet, et al., 2023; Rouchidane Eyitayo, Giraud, et al., 2023). Other teams have described the reconstitution of Bcl-xL in co-formation experiments: Bcl-xL solubilized in detergent is mixed with cholate-solubilized phospholipid and MSP; subsequent detergent removal results in the assembly of the ND and in membrane incorporation of Bcl-xL C-terminal domain; of note, in that protocol, Bcl-xL undergoes urea denaturation, lyophilization, and detergent solubilization (Yao et al., 2016; Yao & Marassi, 2019) while it is not the case in the protocol described herein.

Structural studies on Bak have been carried out on the protein truncated from its C-terminal hydrophobic domain (Moldoveanu et al., 2006). Likewise, many functional studies have used BakΔC to measure its affinity for BH3-only proteins (i.e., Bim, Bad, or PUMA). To this day, only two publications report the use of NDs for Bak studies; Sperl et al. report a high-resolution structural analysis using an engineered version of Bak where the hydrophobic C-terminal domain was replaced by a His6-Tag to allow stable membrane binding on Ni^2+^-caged lipids in MSP-encircled NDs (Sperl et al., 2021); Dadsena instead used SMA Lipid Particles to study what lipids surround BAK during mitochondrial outer membrane permeabilization (Dadsena et al., 2024). Therefore the contribution of the membrane environment on Bak conformational changes is incomplete. In this study we describe a protocol to produce Bak from CFPS, in the presence of NDs assembled from a phospholipid blend mimicking the mitochondrial outer membrane (Asciolla et al., 2012). We describe the purification of ND-inserted Bak; we show that this setup is useful to compare the structural plasticity of Bax and Bak; we describe the spontaneous insertion of Bak into NDs and document the supramolecular assemblies it forms when embedded in the ND bilayer.

## 2. Materials

### 2.1 Common disposables

- TipOne® 1000µL Graduated Tip (#S1111-6000, StarLab)
- TipOne® 200µL Ultra Point Graduated Tip (#S1113-1000, StarLab)
- TipOne® 10µL Graduated Tip (#S1111-3000, StarLab)
- Clearline PP microtube 2 ml, round bottom, snap cap, non sterile (#390692, ClearLine)
- Clearline PP microtube 1.5 ml, conical bottom, snap cap, non sterile (#390690, ClearLine)
- FlipTube® A (#235454, Hamilton)
- Cap locks 1.5ml – 2ml tubes (#I1415-1508, StarLab)
- Spectra/Por® 7 Dialysis membrane, Pre-treated RC tubing, MWCO 10 kDa, (#132118, SpectrumLabs)
- Ni-NTA Superflow (#30410, Qiagen)
- HisTrap FF 1 mL (#17-5319-01, Cytiva)
- Terumo^TM^ Syringe 1 ml luer slip tip (#MDSS01SE, Terumo®)
- Agani^TM^ Needle 20Gx1” (0.9×25mm) (#AN*2025R1, Terumo®)
- Slide-A-Lyzer® Dialysis Cassette MWCO 10kDa 0.1-0.5 ml (#66383, Thermo Scientific)
- Slide-A-Lyzer® Buoys (#66430, Thermo Scientific)
- Mini-PROTEAN TGX Gels 4-15% (#4561083, Bio-Rad)
- NativePAGE^TM^ 4-16% Bis-Tris Gel (#BN2111BX10, Invitrogen)
- Uvette® (#0030106.318, Eppendorf)
- Parafilm® M (#HS234526C, Sigma)
- Vivaspin® 500, MWCO 30 kDa, Polyethersulfone (#28-9322-35, Cytiva)
- Immersol™ 518 F fluorescence-free immersion oil, 20 mL (Cat. No. 444960-0000-000, ZEISS)
- Lens Cleaning Tissue, Grade 105, 100 × 150 mm (Cat. No. 2105-841, Whatman/Cytiva)
- MassGlass™ UC – Sample Prep Kit (MP-CON-21022, Refeyn)
- MassFerence™ P1 Calibrant (MP-CON-41033, Refeyn)
- MP-Alignment Kit (MP-CON-21019, Refeyn)

### 2.2 Cells and reagents

- BL21 (DE3 star) competent E. coli strain (ideal for routine T7 expression, with reduced protease activity to increase protein production yield and stability)
- TRIS base (#200923-A, Euromedex)
- Sodium chloride (#1112-A, Euromedex)
- Imidazole (#I202, Sigma-Aldrich)
- Sodium cholate hydrate (#C1254, Sigma-Aldrich)
- Sodium carbonate anhydrous (#27771.290, VWR)
- â-mercaptoethanol (#M6250, Sigma-Aldrich)
- Phospholipids : POPC (16:0-18:1 PC • 1-Palmitoyl-2-Oleoyl-sn-Glycero-3-Phosphocholine, #P516, Anatrace), POPE (16:0-18:1 PE • 1-Hexadecanoyl-2-(9Z-Octadecenoyl)-sn-Glycero-3-Phosphoethanolamine • 1-Palmitoyl-2-Oleoyl-sn-Glycero-3-Phosphoethanolamine, #P416, Anatrace), DOPS (18:1 PS • 1,2-di-(9Z-octadecenoyl)-sn-glycero-3-phospho-L-serine (sodium salt) • 1,2-Dioleoyl-sn-glycero-3-PS • 1,2-dioleoyl-sn-glycero-3-phosphatidylserine, #D718, Anatrace), Cardiolipin (#710335P, Sigma-Aldrich)
- Plasmids : pIVEX2.3-Bcl-xL (human Bcl-xL was cloned between NdeI-XhoI, and mutagenesis was used to insert a second STOP codon after the first one, to prevent unwanted readthrough), pIVEX2.3-Bax, pIVEX2.3-Bak (human Bak was cloned between NdeI-SmaI sites). pIVEX2.3-His6Bak was a kind gift from Lucie Bergdoll. pET28a-MSP1E3D1 (#20066, Addgene).
- All the regents for cell-free production in continuous exchange are listed in the original protocol (Schwarz et al., 2007) except RNasin® Ribonuclease Inhibitor (N2511, Promega) is used to prevent RNA degradation.
- Antibodies: Rabbit monoclonal anti-Bcl-xL (#ab32370, Abcam) 1/4000 dilution, mouse monoclonal anti-human Bax 2D2 (Santa-Cruz) 1/5000 dilution, rabbit polyclonal Bak NT (#06-536, Millipore) 1/4000 dilution, mouse monoclonal anti-his tag (clone 4E3D10H2/E3, Thermo Fisher) 1/5000 dilution. Peroxidase-AffiniPure Goat Anti-Rabbit IgG (H+L) (#111-035-045, Jackson ImmunoResearch) 1/10,000. Peroxidase AffiniPure Goat Anti-Mouse IgG (H+L) (#115-035-003, Jackson ImmunoResearch) 1/10,000.
- MFP1 proteins standards (Refeyn, MP-CON-41033)
- BSA protein (Sigma, A2153)

### 2.3 Equipment

- Ultrasonic water bath Elmasonic Facile 30 h (#1117697001, Elma)
- Temperature-controlled shaker for cell-free production at 28°C overnight under gentle agitation
- ÄKTA chromatography system for protein purification
- Superdex 200 Increase 10/300 GL (Cytiva)
- DynaPro NanoStar (Wyatt Technology) for Dynamic Light Scattering (DLS) analyses, to measure the size distribution and homogeneity of macromolecules, and determine their hydrodynamics radius
- TwoMP (Refeyn) for Mass Photometry (MP) analyses, to determine the molecular mass, oligomeric state, sample heterogeneity, and relative abundance of biomolecular species in solution at the single-molecule level

### 2.4 Software

ImageJ for densitometric analyses of images acquired from western blotting experiments. Prism v. 8 (Graph Pad) for statistical analyses and data presentation.

AcquireMP v2025 R1.2 (Refeyn) for acquisition and recording of Mass Photometry (MP) measurements. DiscoverMP v2026 R1.1 (Refeyn) for processing, analysis, and interpretation of Mass Photometry (MP) data.

## 3. Methods

### 3.1 Production of nanodiscs (NDs)

Membrane-inserted Bak is produced using home-made NDs. The composition and proportions of phospholipids (see 3.1.2) are chosen to mimic the outer mitochondrial membrane of animal cells (48% POPC, 32% POPE, 12% DOPS, 8% cardiolipin). The membrane scaffold protein encircling phospholipid bilayer is MSP1E3D1, with an N-terminal His7-tag to enable affinity-purification of NDs. Like all MSP proteins, this variant was genetically engineered based on the sequence of Apo-A1 devoid of its globular N-terminal domain. Specific to this variant are the removal of the first 11 amino acids in the Helix 1, and the repeat of Helix 4, 5 and 6 between Helix 6 and Helix 7. The resulting NDs diameter typically ranges between 8 and 15 nm.

#### 3.1.1 Production and purification of His7-MSP1E3D1

The protocol to produce and purify the scaffold protein His7-MSP1E3D1 is adapted from one of our former publications (Rouchidane Eyitayo, Giraud, et al., 2023). The plasmid encoding His7-MSP1E3D1 is transformed into competent E. coli BL21(DE3*) and an isolated clone is grown overnight at 37°C under agitation in 25 mL of LB medium. The culture is further transferred in 500 mL of TB medium (37°C under agitation) until DO_600nm_=0.8. Protein expression is induced with 1 mM IPTG for 3 hours. Cells are centrifuged and lysed by sonication. After a centrifugation at 46,000 x *g* for 30 min at 4°C, cell debris are discarded and the supernatant is loaded for at least 4h at 4°C onto a 2 mL Ni-NTA column in a closed circuit at 1 mL/min. The column is washed successively with 5 column volumes (CV) of 40 mM Tris-HCl (pH 8.0), 300 mM NaCl, 1% Triton X-100, followed by 5 CV of 40 mM Tris-HCl (pH 8.0), 300 mM NaCl, 50 mM sodium cholate. The column is then connected to an ÄKTA purifier (see Notes for ÄKTA HisTrap chromatography parameters) and washed with 5 CV of buffer A (40 mM Tris-HCl (pH 8.0), 300 mM NaCl). The protein is finally eluted in a two-step process with first, 7% Buffer B (40 mM Tris-HCl (pH 8.0), 300 mM NaCl, 300 mM imidazole) followed by 100% of buffer B. The eluate is dialyzed in 10 mM Tris-HCl (pH 8.0), 100 mM NaCl, and concentrated to 7 mg/mL. Working aliquots are stored at -80°C.

#### 3.1.2 ND production and purification

NDs are produced and purified with adaptations from one of our previous publications (Rouchidane Eyitayo, Giraud, et al., 2023). The step-by-step procedure is the following.

1. Lipids stock solutions are stored in chloroform at 25 mg/mL. A 5 mg mitochondria-mimicking lipid mixture is assembled from 2.4 mg of POPC, 1.6 mg of POPE, 0.6 mg of DOPS and 0.4 mg of cardiolipin. The mixture is dried under Argon or Nitrogen flux for 1 to 3 hours to ensure chloroform evaporation.
2. The lipid mixture is resuspended in 140 µL of 6 mM Tris (pH 8.0), 60 mM NaCl, 200mM sodium cholate.
3. This solution is sonicated for 10 min (discontinuous pulse) in an ultrasonic water bath.
4. 3 mg of purified His7-MSP1E3D1 are mixed with the 5 mg lipid mixture, and incubated for 1h at room temperature on a rotating wheel.
5. A first dialysis is performed when the lipid-cholate mix is transferred into a Spectra/Por 7 dialysis membrane tubing (MWCO 10 kDa) and immersed in 500 mL of 10 mM Tris (pH 8.0), 100 mM NaCl at room temperature for 2 hours at least. A second identical dialysis is performed overnight at 4°C.
6. The dialized mixture is centrifuged 20,000 x *g* for 15 min at room temperature, and the supernatant recovered. At this step, it is possible to concentrate the sample to decrease its volume before injection onto the ÄKTA chromatography system. The sample is always filtrated at 0,2µm before loading onto ÄKTA.
7. NDs are purified by a size exclusion chromatography (SEC) on a Superdex 200 10/300 GL increase column (see Notes for ÄKTA Superdex 200 10/300 GL increase chromatography parameters).
8. SEC chromatogram are analyzed: typically, NDs elute in 2 consecutive but separate peaks (Figure 1), the respective relative intensity of which may vary between experiments. 500 µL fractions corresponding to the peaks are collected, and further processed individually on a DynaPro NanoStar (see Notes for DLS parameters). DLS analyses enable the selection of SEC fractions containing monodisperse macromolecules, while the rest is discarded. Then the fractions containing NDs of comparable hydrodynamics radius are pooled and concentrated with Vivaspin tubes (MWCO 30 kDa) to about 200 µL. The protein content is finally determined using Pierce BCA Protein Assay Kit and ideally ranges between 60-80 µM.

**Figure 1.**
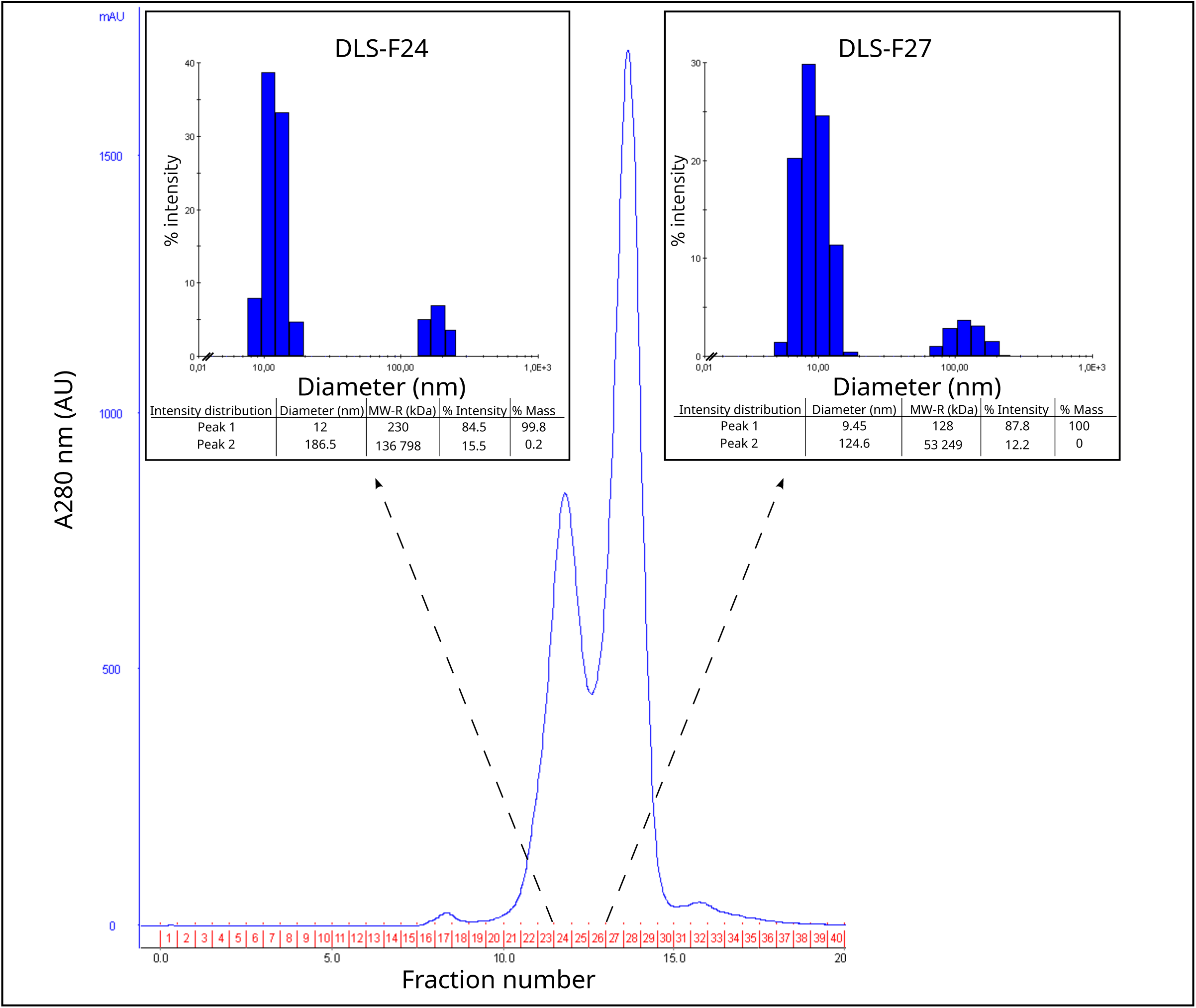
-Quality control of His7-MSP1E3D1 NDs. Size exclusion chromatogram (Superdex Increase 200 10/300 GL column) show absorbance recordings at 280 nm in the fractions collected. Each fraction of interest was analyzed by DLS (examples are shown in inserts for fractions 24 and 27) to collect information on NDs diameter, % intensity and % mass. For each fraction, a triplicate of 10 measures was acquired, and the mean of each triplicate was generated. Inserts are representative of 1 mean. Following DLS, fractions 24-25-26 were pooled for peak 1 and 27-28-29 for peak 2. After concentration, protein content was determined and NDs were stored at 4°C.

### 3.2 Production of Bcl-2 family proteins by CFPS with NDs

Untagged full-length human Bcl-xL and Bax, and His6-Bax are expressed as previously described (Rouchidane Eyitayo, Boudier-Lemosquet, et al., 2023; Rouchidane Eyitayo, Giraud, et al., 2023), while Bak, depending on the experiments, is either untagged or harbors a His6-tag in N-term. Bak truncated from its C-terminal hydrophobic helix (BakΔC) was obtained by introducing a stop codon in position 182 to exclude the last 30 amino-acids.

Cell-free protein synthesis is performed as initially described (Rouchidane Eyitayo, Giraud, et al., 2023): the step-by-step procedure is as follows.

1. 1.5 mL Flip-Tube A are chosen because their cap is best fitted to hold the 100 µL of Reaction Mix. Scissors are used to detach the cap, and to cut off the conical bottom of the Flip-Tube A.
2. The Reaction Mix containing the plasmid of interest and 3.5 µM of NDs is carefully deposited into the inverted cap.
3. This mixture is covered with a 1.5 cm x 1.5 cm single-layered square cut from a dialysis membrane tubing (MWCO 10 kDa). The tube is reassembled upside down and the dialysis membrane is secured in place by a plastic clip to seal closed the cap and tube.
4. Through the open end of the Flip-Tube A, 1700 μL of Feeding Mix are added, and the opening is sealed with Parafilm.
5. The setup is incubated at 28°C under gentle agitation (80 rpm) overnight.

#### 3.2.1 Reaction Mix recovery and production analysis

##### IMPORTANT NOTE

For every analysis step (production and affinity chromatography), the volumes of the fractions are estimated on a precision scale. Therefore, the microfuge tubes used to receive the fractions must be weighted empty first, and once loaded with the fractions. The difference indicates the fraction volume, and the same percentage of pellet/supernatant or unbound/eluate is separated onto a SDS-PAGE for comparative analyses.

1. The Feeding Mix is discarded with a P1000 pipette. The Reaction Mix is carefully harvested from the cap with a P200 pipette : the tip pierces the dialysis membrane, and the volume is transferred into a new microfuge tube. A centrifugation of 15 min at 20,000 x *g* and 4°C separates insoluble proteins (pellet) from soluble proteins (supernatant).
2. The supernatant S1 is weighted on a precision scale. A 5% aliquot is taken for western blotting analysis.
3. The pellet P1 is washed with 100 µL of 25 mM Tris (pH 8.0), 250 mM NaCl and centrifuged again for 15 min at 20,000 x *g* and 4°C. The wash is discarded and the pellet is resuspended in SDS-PAGE 2x sample buffer (125 mM Tris-HCl (pH 6.8), 4% SDS, 20% Glycerol and bromophenol blue), with the same volume as that previously determined for S1.
4. S1 and P1 are finally analyzed by SDS-PAGE followed by western blotting to detect on one gel the protein produced, and on another gel His7-MSP1E3D1. The same percentage of S1 and P1 fractions is analyzed. This percentage depends on the protein production yield : typical percentage for Bcl-xL is 0.01%, for Bax is 0.05%, and for Bak is 0.5%. For anti-His immunodetection, 0.5% is analyzed.

#### 3.2.2 Affinity purification of NDs and associated/embedded proteins

1. Following production, S1 is diluted 1:1 with binding buffer (25 mM Tris (pH 8.0), 250 mM NaCl) and incubated with 60 μL of Ni-NTA Sepharose slurry (previously equilibrated with binding buffer) for 2 h at 4°C on a rotating wheel.
2. After centrifugation of 1 min at 1,000 x *g* at room temperature, the unbound fraction U1 is collected and its volume estimated on a precision scale. A 5% aliquot is taken for western blotting analysis and the remaining fraction stored at -20°C.
3. The resin is washed 3 times with 500 µL of washing buffer (25 mM Tris (pH 8.0), 250 mM NaCl, 20 mM Imidazole).
4. A volume of elution buffer (25 mM Tris (pH 8.0), 250 mM NaCl, 300 mM Imidazole (pH 8.0)) equal to that of U1 is added to the resin and incubated for 5 min on ice. After a spin of 1 min at 1,000 x *g* at room temperature, the eluate fraction E1 is collected and its actual volume is estimated on a precision scale. A 5% aliquot is taken for western blotting analysis.
5. U1 and E1 are finally analyzed by SDS-PAGE followed by western blotting to detect on one gel the protein produced, and on another gel HIS7-MSP1E3D1. The same percentage of U1 and E1 fractions is analyzed. This percentage depends on the protein production yield : typical percentage for Bcl-xL is 0.01%, for Bax is 0.05%, and for Bak is 0.5%. For anti-His immunodetection, 0.5% is analyzed.

As a quality indicator of the experiment, NDs should only be detected in the E1 fraction, with no His signal detected at all in the U1 fraction. Granted this, the distribution of the Bcl-2 proteins produced may be analyzed to determine what proportion did not bind to NDs (Unbound), and what proportion was associated to/inserted into NDs (Eluate). Image J is used to proceed to a densitometric analysis of the signals detected in U and E.

#### 3.2.3 Purification of ND-embedded proteins

To distinguish between proteins that are associated to NDs by electrostatic interactions, and those inserted into NDs, a sodium carbonate treatment (Na_2_CO_3_^-^) is performed. The kosmotropic agent disrupts the hydrogen bonding network and salt bridges established between amino acids and the polar heads of the phospholipids, while the hydrophobic interactions established after insertion of protein transmembrane domains are retained. After this treatment, a second Ni-NTA affinity purification step is performed. “Associated proteins” are collected in the unbound fraction U2, while “ND-embedded proteins” are recovered in the eluate fraction E2.

Before proceeding to the carbonate treatment, imidazole must be dialyzed from E1.

1. E1 fraction is transferred from a syringe equipped with a needle into a Slide-A-Lyzer dialysis cassette (MWCO 10 kDa). Dialysis is carried out in 500 mL of binding buffer for 2h at 4°C, and repeated once, over night. Alternatively, dialysis can be repeated twice for 2h at 4°C.
2. E1 is recovered with a syringe equipped with a needle, and the volume estimated on a precision scale. The sample is divided in two. One tube is added with 100 mM Na_2_CO_3_^-^, the second tube is instead added with an equal volume of binding buffer. The mix is incubated for 15 min on ice, and then incubated with 60 μL of Ni-NTA Sepharose slurry (previously equilibrated in binding buffer) for 2 h at 4°C on a rotating wheel.
3. Affinity chromatography proceeds as described in section 3.2.2. The fractions collected are U2 and E2 of Na_2_CO_3_^-^ treated samples, and U2 and E2 of untreated samples. All the fractions are finally analyzed by SDS-PAGE followed by western blotting to detect the protein(s) produced, and in parallel His7-MSP1E3D1 as a quality control of the affinity chromatography. The same percentage of U2 and E2 fractions is analyzed. This percentage depends on the protein production yield : typical percentage for Bcl-xL is 0.5%, for Bax is 0.2%, and for Bak is 1.5%. For anti-His immunodetection, 1% is analyzed.

As a quality indicator of the experiment, NDs should only be detected in the E2 fraction, with no His signal detected at all in the U2 fraction. Granted this, the proportion of ND-inserted Bcl-2 proteins can be appreciated in the E2 fraction compared to proteins removed in the U2 fraction. Image J is used to proceed to a densitometric analysis of the signals detected in U and E.

This protocol is very efficient to purify ND-inserted Bcl-2 proteins, but it is important to bear in mind that it does not allow to discard empty NDs. Hence, an optimization step might be necessary to facilitate subsequent structure/function analyses. Notably, screening a range of ND concentrations in the Reaction Mix might allow to decrease the proportion of empty NDs.

#### 3.2.4 Supramolecular assembly of Bak in NDs

In biological membranes, Bak oligomerizes and creates pores in the mitochondrial external membrane to release apoptogenic factors essential for the final steps of apoptosis. Whether artificial membranes organized in NDs are a good proxy to allow Bak oligomerization and facilitate the structural study of such supramolecular assemblies was the focus of this study. Several approaches, providing complementary information, have been used.

- Human Bak harbors 2 cysteine residues that might be conducive to cysteine-dependent dimers. The formation of such dimers was evaluated by loading E1 fraction on gradient SDS-PAGE and comparing samples incubated with and without β-mercaptoethanol.
- The arrangement of Bak molecules inserted in close proximity to one another in the ND bilayer can be probed by cross-linking experiments; the length of the chemical cross-linker sets the distance between molecules in adducts identified on denaturing gradient gels. For these experiments, E1 fraction (containing NDs and associated proteins) is dialyzed at 4°C in 10 mM HEPES (pH 7.4), 100 mM NaCl. Samples are either mock-treated or incubated with 0.01% Brij 58 at room temperature for 15 minutes to solubilize NDs. Proteins within 7.7 Å are then cross-linked with 0.2 mM of disuccinimidylglutarate (DSG) for 30 min at room temperature. The reaction is finally quenched with 50 mM of Tris-base for 15 min at room temperature. Samples are diluted 1:1 with 2x loading buffer prior to separation on 4-16% Mini-Protean TGX Precast PAGE gels (Bio-Rad) and immunodetection of Bak and His7-MSP1E3D1. Adducts in mock conditions represent complexes formed in NDs, whereas Brij 58-treated samples reveal complexes that survived solubilization.
- The supramolecular organization of Bak in NDs can also be analyzed on native gels. In that case, E1 fraction is analyzed on a non-denaturing Bis-Tris 4%-16% gradient gel following the procedure described in (Dewson et al., 2008). Briefly, protein separation is performed at 150 V for ∼1.5 hours. Gels are further incubated with transfer buffer (25 mM Tris, 192 mM Glycine, 20% Ethanol) supplemented with 0.1% SDS for 30 min under gentle agitation at room temperature. Proteins are transferred onto PVDF membranes at 30 V, overnight at 4°C. The membranes are then incubated in the migration buffer (25 mM Tris, 192 mM Glycine, 1% SDS) supplemented with 5% SDS and 2% β-mercaptoethanol for 30 min at 65°C under gentle agitation. After several washes with transfer buffer (without ethanol) to eliminate SDS and β-mercaptoethanol, the standard immunodetection procedure is carried out.
- Finally, mass photometry (MP) is an emerging label-free method for the quality control of supramolecular assemblies en route for high-resolution structural determination. By measuring the molecular mass and deducing stoichiometry, it enables the characterization of proteins reconstituted in NDs. It discriminates properly assembled complexes from contaminants, and assesses aggregates and sample homogeneity. This method helps optimizing sample quality before cryo-electron microscopy experiments. Sample preparation was as follows: empty ND were obtained as described in section 3.1.2. Bak in NDs correspond to eluate E1 of Histrap 1 (section 4.2) which was dialyzed as described in section 3.2.3. Mass photometry measurements were performed with a Two MP instrument (Refeyn) in a well on a glass slide according to the supplier’s instructions. Filtered buffers, samples and calibration standards were equilibrated at room temperature before use. Focus of the objective was done with 10 μL of reference buffer 25 mM Tris (pH 8.0), 250 mM NaCl. Then, 10 μL of sample at a concentration of 20 nM were added to the drop, mixed by pipetting up and down 3–4 times, to obtain a final concentration of 10 nM. The acquisition software AcquireMP recorded data for 1 min. Analysis was performed using DiscoverMP software. Calibration was done with MFP1 proteins standards (Refeyn, MP-CON-41033) and with BSA protein (Sigma, A2153).

## 4. Results and analysis

### 4.1 Production of Bcl-xL, Bax and Bak by CFPS

#### 4.1.1 Comparison of Bcl-xL, Bax and Bak production yields and solubility

Our team already described the use of the current CFPS protocol to produce human Bcl-xL and Bax(Rouchidane Eyitayo, Boudier-Lemosquet, et al., 2023; Rouchidane Eyitayo, Giraud, et al., 2023). We now extend it to human Bak. To compare respective production yields, the three proteins were synthesized separately overnight (section 3.2), and the reaction mixes recovered were centrifuged at 20,000 x *g*. The supernatant contains soluble proteins, while insoluble proteins are recovered in the pellet. Figure 2 shows that Bcl-xL (Fig. 2A), Bax (Fig. 2B) and Bak (Fig. 2C) are recovered in supernatant, with similar solubility (∼100%) calculated from independent western blots (Fig. 2D). Yet, Coomassie Blue staining of 1% of each production provides a direct indication that Bcl-xL (Fig. 2A) and Bax (Fig. 2B) are well produced (arrows) while Bak relative abundance is less (Fig. 2C) since its detection requires immunoblotting.

**Figure 2.**
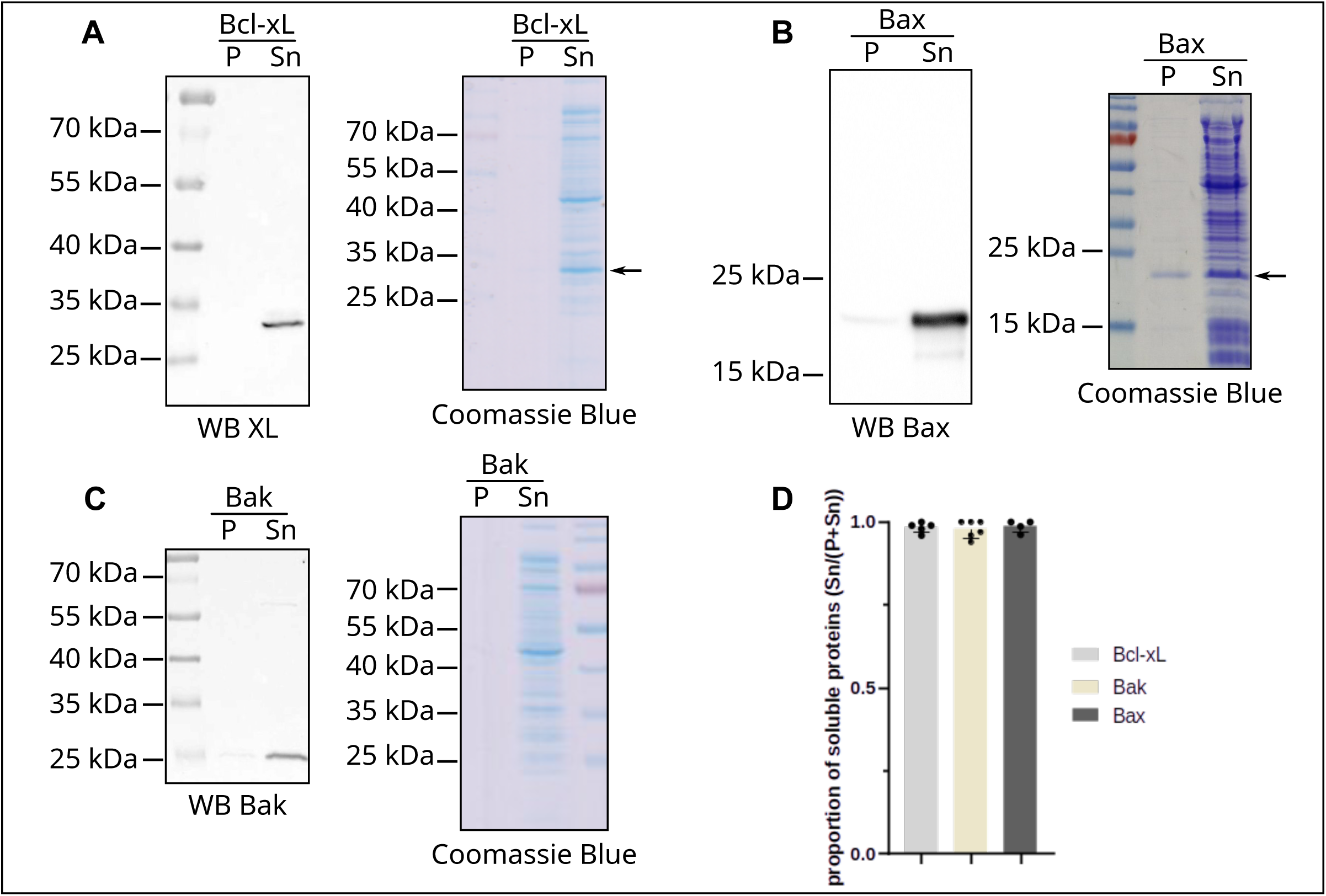
-Production yields and solubilty comparisons for CFPS of human Bcl-xL, Bax and Bak. Bcl-xL, Bax or Bak (respectively A, B and C) were produced from cell-free bacterial extracts. Fractions were separated on SDS-PAGE and analyzed by western blot (left) and Coomassie Blue staining (right). 1% of Pellet (P) or Supernatant (Sn) were analyzed by Coomassie Blue staining. For western blot analysis, 0.01% of Bcl-xL and Bax fractions were analyzed, and 0.5% for Bak. D-Densitometric quantification of n>3 independent experiments. Mean was calculated +/-SD, and one-way ANOVA test showed no statistical difference.

#### 4.1.2 Comparison of Bax and Bak for the effects of N-terminal His-tag and ND on their solubility in CFPS

Bcl-2 family proteins exhibit remarkable structural plasticity: conformational changes allow accommodations needed for dynamic interactions and/or higher order assemblies, and in the case of multi-domain pro-apoptotic proteins, they account for stepwise functional activation. The sequential conformational changes of Bax were initially documented in cells, when the protein was expressed as an N-or C-terminal fusion with affinity tags (Cartron et al., 2003, 2005); more recently, CFPS proved a converging technique to identify NDs as a catalytic trigger to prime the mobility of Bax N-terminus (in the absence of tag): indeed we reported that, while Bax expressed by CFPS is soluble, the addition of NDs to the Reaction Mix results in a dramatic 50% drop in Bax solubility. The molecular determinants of this intrinsic mobility of Bax N-terminus were further characterized in (Rouchidane Eyitayo et al., 2024). Strikingly, CFPS further revealed that the addition of an N-terminal His6-tag was not a primary trigger for Bax structural activation, as the tagged protein exhibits a solubility comparable to the untagged one (Fig. 3A&B). This trait applied both in the absence and in the presence of ND, underscoring that the triggering effect of lipids prevails over the relative instability imparted by the N-terminal His tag.

**Figure 3.**
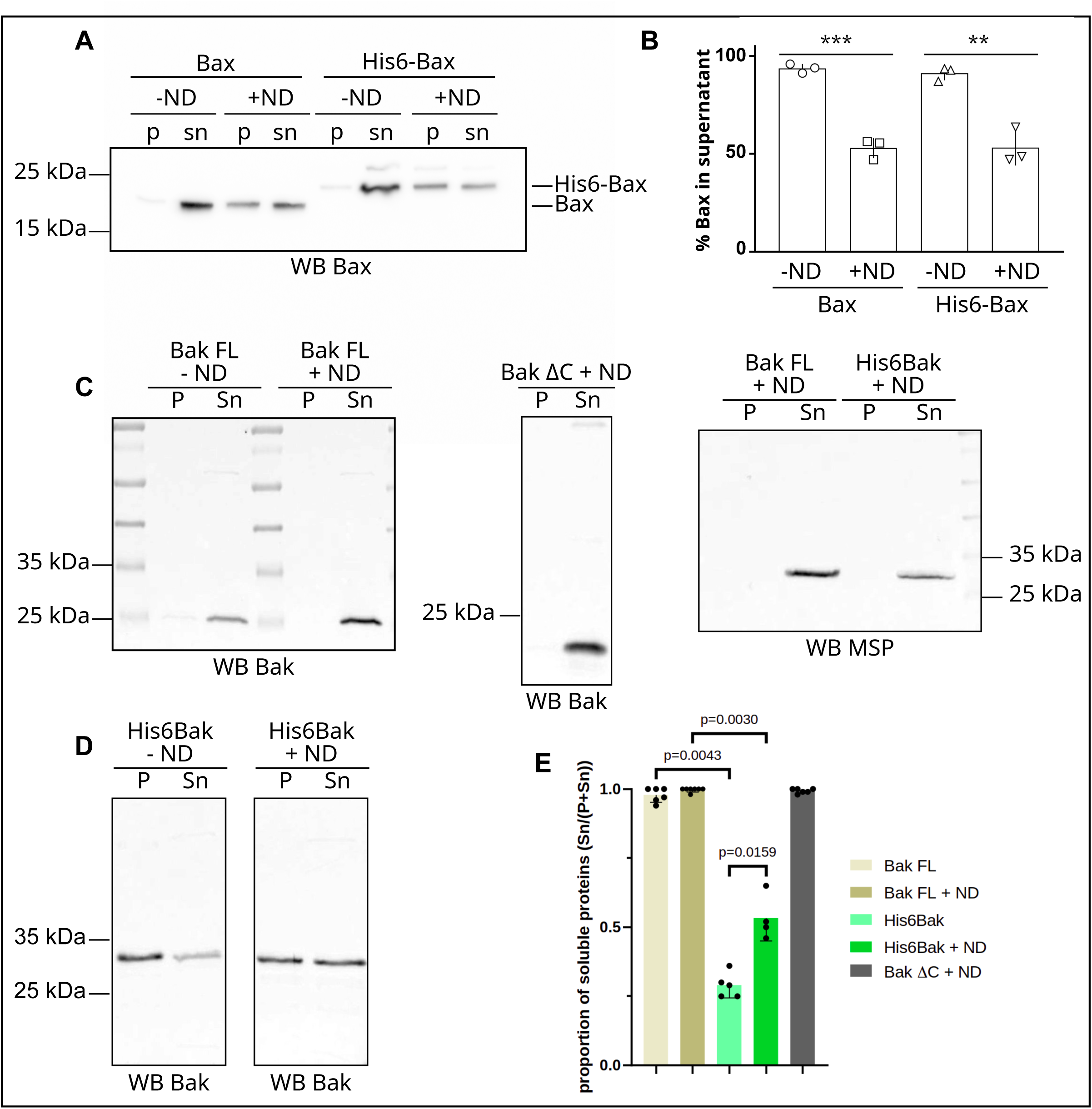
-Effect of NDs and N-term His6 tag on the solubility of Bak compared to Bax. The indicated proteins were produced by CFPS without NDs or added with NDs. After the 20,000 x *g* centrifugation, the same proportion of Pellet (P) and Supernatant (S) fractions were separated on SDS-PAGE followed by immunodectection. A-Comparison of Full-Length-Bax and His6-Bax. NDs are 1.5 µM, 0.1% of the fractions is analyzed. B-Densitometric quantification of 3 independent experiments. Mean is shown +/-SD with a t-test (unpaired, parametric): *** p<0.001; ** p<0.01. C-Comparison of Full-Length Bak (Bak FL) to C-terminal truncated Bak (BakΔC) or to His6-Bak (D-). NDs are 3.5 µM. 0.5% of the fractions were separated by SDS-PAGE followed by immunodetection of Bak. E-Densitometric quantification analysis of n>3 independent experiment. Mean is shown +/-SD, with a statistical Mann-Whitney test.

We further explored how the knowledge earned from Bax transposes to Bak. After NDs have been produced and purified as described in Fig. 1 (section 3.1.2), CFPS of Bak was achieved in the presence or in the absence of NDs, and the Reaction Mixes were centrifuged. Full Length Bak (Bak FL) was 100% soluble regardless of the presence of NDs (Fig. 3C), thus mirroring BakΔC truncated of its C-terminal transmembrane domain, which was used as a control. Surprisingly, adding an N-terminal His6-tag to Bak entailed a dramatic loss of solubility, as ∼70% of the protein was recovered in the pellet fraction (Figure 3D&E), while NDs (i. e. His7-MSP) were consistently detected in the soluble fraction (Fig. 3C). Therefore, unlike Bax, the N-term His6-tag sufficed to unlock Bak conformational change. Moreover, the presence of NDs during CFPS partially restored His6-Bak solubility to 50% (Figure 3D&E). This set of experiments thus highlighted striking differences between Bax and Bak structural plasticity, and the respective triggering effects played by NDs for the former, and N-term tag for the latter, to prime conformational changes.

### 4.2 Association of Bak to NDs

Untagged Bak FL, recovered in the Reaction Mix after overnight CFPS, qualifies as a fully soluble protein after centrifugation. Since NDs co-fractionate in the supernatant, we further addressed the ability of Bak to physically associate with NDs. The NDs produced according to the protocol herein (section 3.1) are encircled by His7-MSP1E3D1 proteins. The procedure for Ni-NTA affinity purification described in section 3.2.2 was followed to assay Bak association to NDs. Separation on SDS-PAGE of the eluate E1 and unbound U1 fractions after Ni-NTA affinity chromatography (Fig. 4A) showed that Bak FL was only recovered in the eluate fraction (quantified in Fig. 4B). On the contrary, BakΔC was found mainly in the unbound fraction (75%), suggesting that Bak association with NDs is driven, for the most part, by its trans-membrane domain.

**Figure 4.**
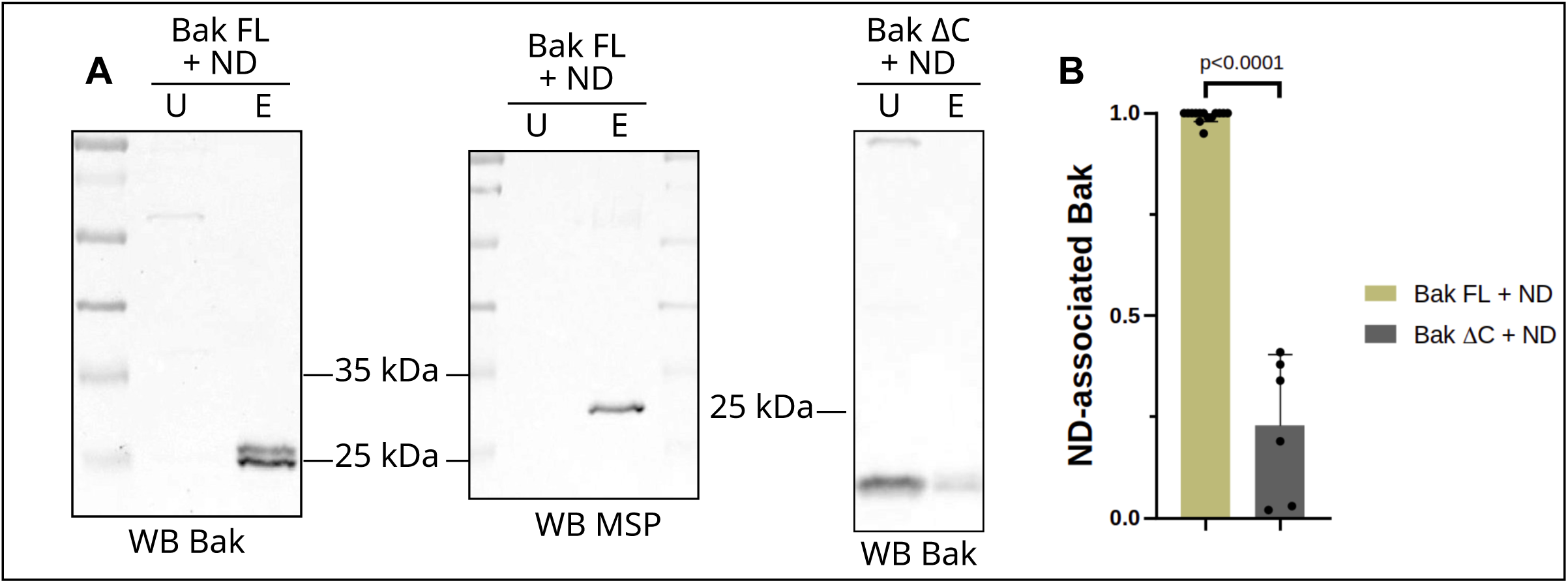
-Association of Bak FL and BakΔC with NDs. A-Bak FL or BakΔC were produced by CFPS in the presence of preformed NDs stabilized by His7-MSP1E3D1. The supernatant was submitted to His-Trap to separate free-soluble Bak (Unbound, U) from ND-associated Bak (Eluate, E). 0.5% of each fraction was separated by SDS-PAGE, followed by immunodetection of Bak. B-Densitometric quantification of n>3 independent experiments. Mean is shown +/-SD, and a Mann Whitney statistical test was used.

### 4.3 Insertion of Bak into NDs

To further discriminate proteins that are associated to NDs by hydrogen bonds and salt bridges, from those inserted into NDs, eluates from the former HisTrap purification were treated with Na_2_CO_3_^-^ and submitted to a second HisTrap chromatography, following protocol from section 3.2.3. We continued the comparison between Bak FL and BakΔC to qualify the type of interaction established by each protein with NDs.

Eluates E2 and unbound U2 fractions post-carbonate or mock treatment were analyzed by SDS-PAGE followed by western blotting (Fig. 5). We observed that the small amount of BakΔC found associated to NDs after HisTrap 1, resisted the carbonate treatment and was again essentially recovered in the E2 fraction. This indicated that BakΔC establishes hydrophobic interactions with NDs. The portion of the protein engaged in these interactions was not investigated further here, but could involve its core. In parallel, we observed that Bak FL was again 100% recovered in the E2 fraction. This indicated that Bak-NDs interaction was not disrupted by the carbonate treatment, and we therefore conclude that Bak was efficiently inserted in the ND bilayer.

**Figure 5.**
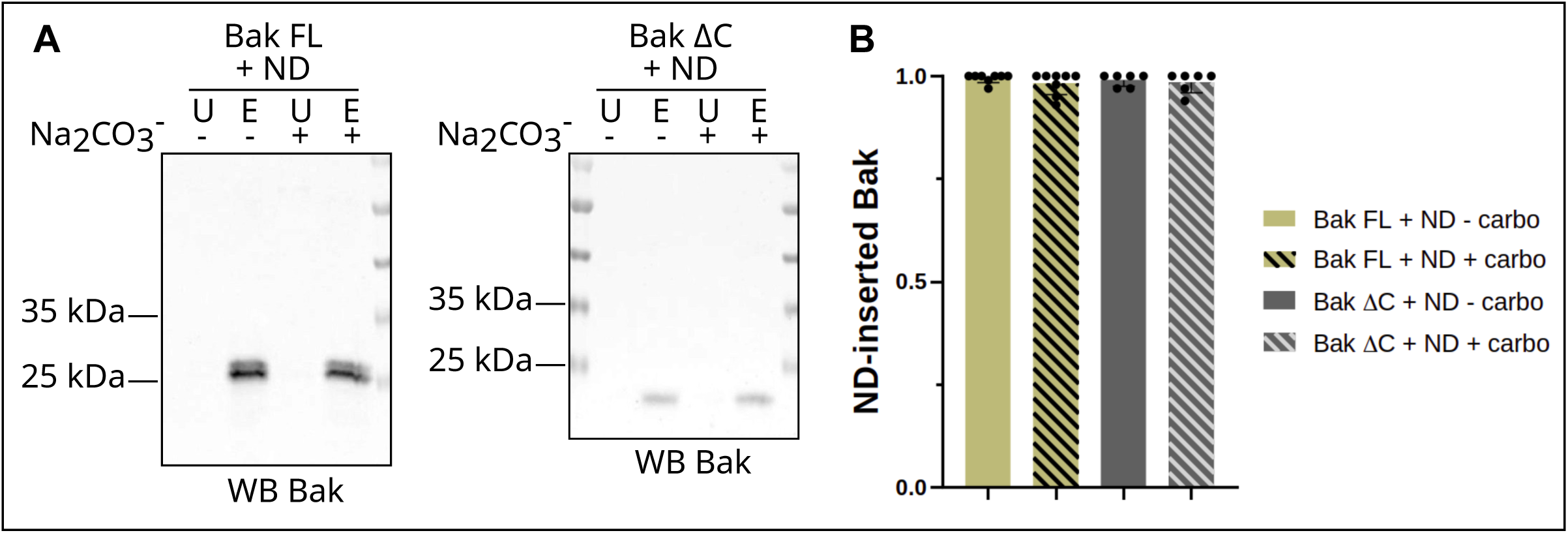
-Insertion of Bak FL and BakΔC into NDs. A-Eluates E1 from the first His-Trap were treated with sodium carbonate and further submitted to a second His-Trap to separate ND-associated proteins (Unbound) from ND-inserted proteins (Eluate). 1,5% of each Bak FL fraction and 5% of each BakΔC fraction were separated by SDS-PAGE, followed by immunodetection of Bak. B-Densitometric quantification of n>3 independent experiments. Mean is shown +/-SD, and a Mann Whitney statistical test was used.

Consequently, this set of experiments demonstrates that the protocol described here allows to produce and purify membrane-inserted full-length Bak.

### 4.4 Supramolecular assemblies of Bak in NDs

The stepwise activation of Bak in the outer membrane of mitochondria proceeds through the sequential formation of higher order assemblies, composed at least of Bak oligomers, and potentially involving other proteins (Cheng et al., 2003; Lalier et al., 2022). Having previously described a protocol for the purification of Bak FL inserted into NDs, we further aimed to assess the supramolecular assemblies and potential oligomeric states of Bak in NDs. The protocol described in section 3.2.4 was followed.

Human Bak sequence contains 2 cysteines that could potentially contribute to the spontaneous formation of cysteine-dependent oligomers. To evaluate this possibility, Bak was synthesized by CFPS in the presence of NDs, and was further purified by NiNTA affinity chromatography. The E1 fraction was incubated with or without β-mercaptoethanol and analyzed on gradient SDS-PAGE. Figure 6A shows that monomeric Bak (23.4 kDa) migrates at ∼25 kDa, and spontaneously produces ∼50 kDa dimers and ∼70 kDa trimers in NDs. These assemblies are resolved when cysteines are reduced by β-mercaptoethanol. The relative intensities obtained from densitometric analyses indicate that ∼50% of Bak remains monomeric in NDs, while the other half arranges mostly in dimers, leaving trimers below 10% (Fig. 6B).

**Figure 6.**
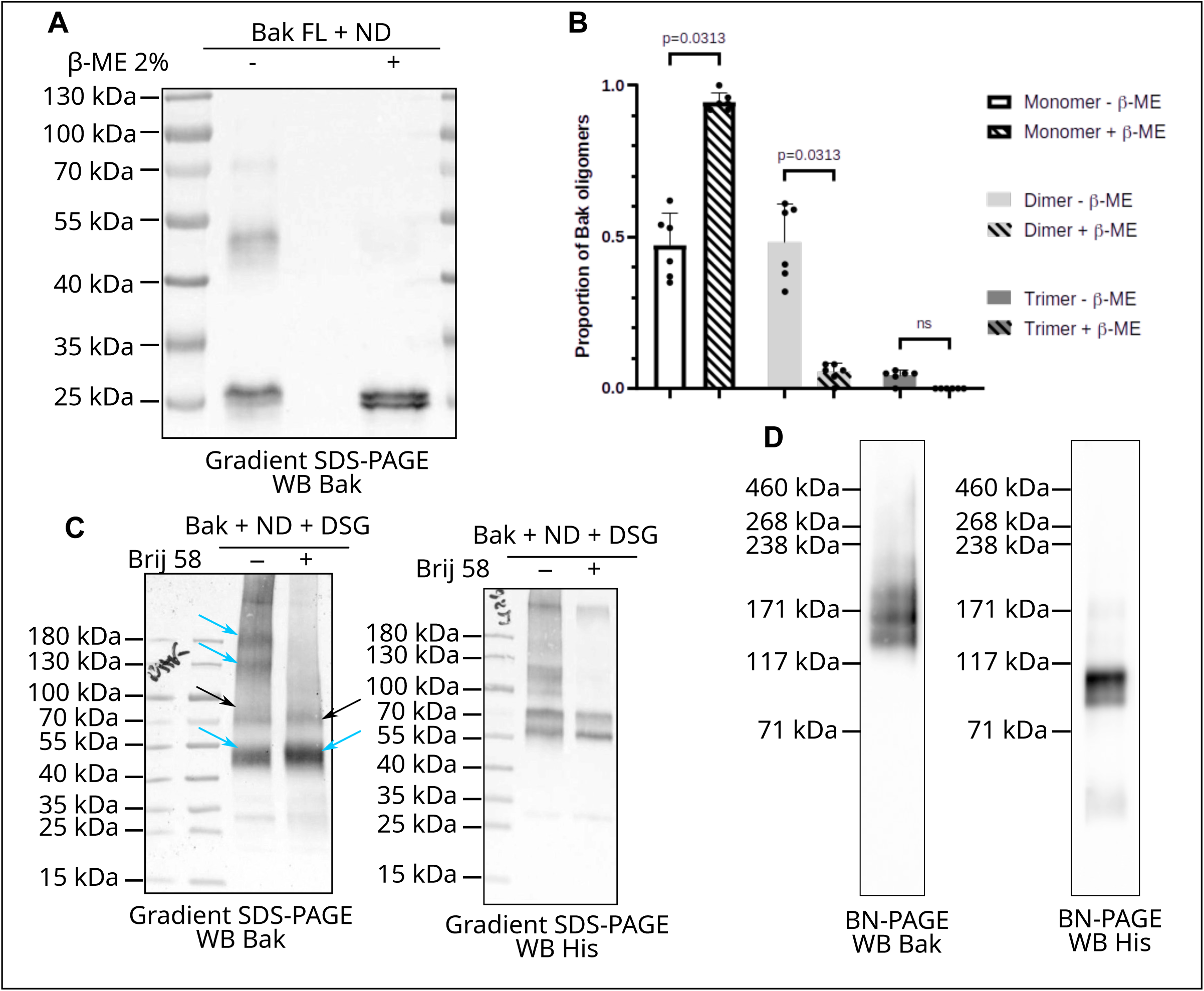
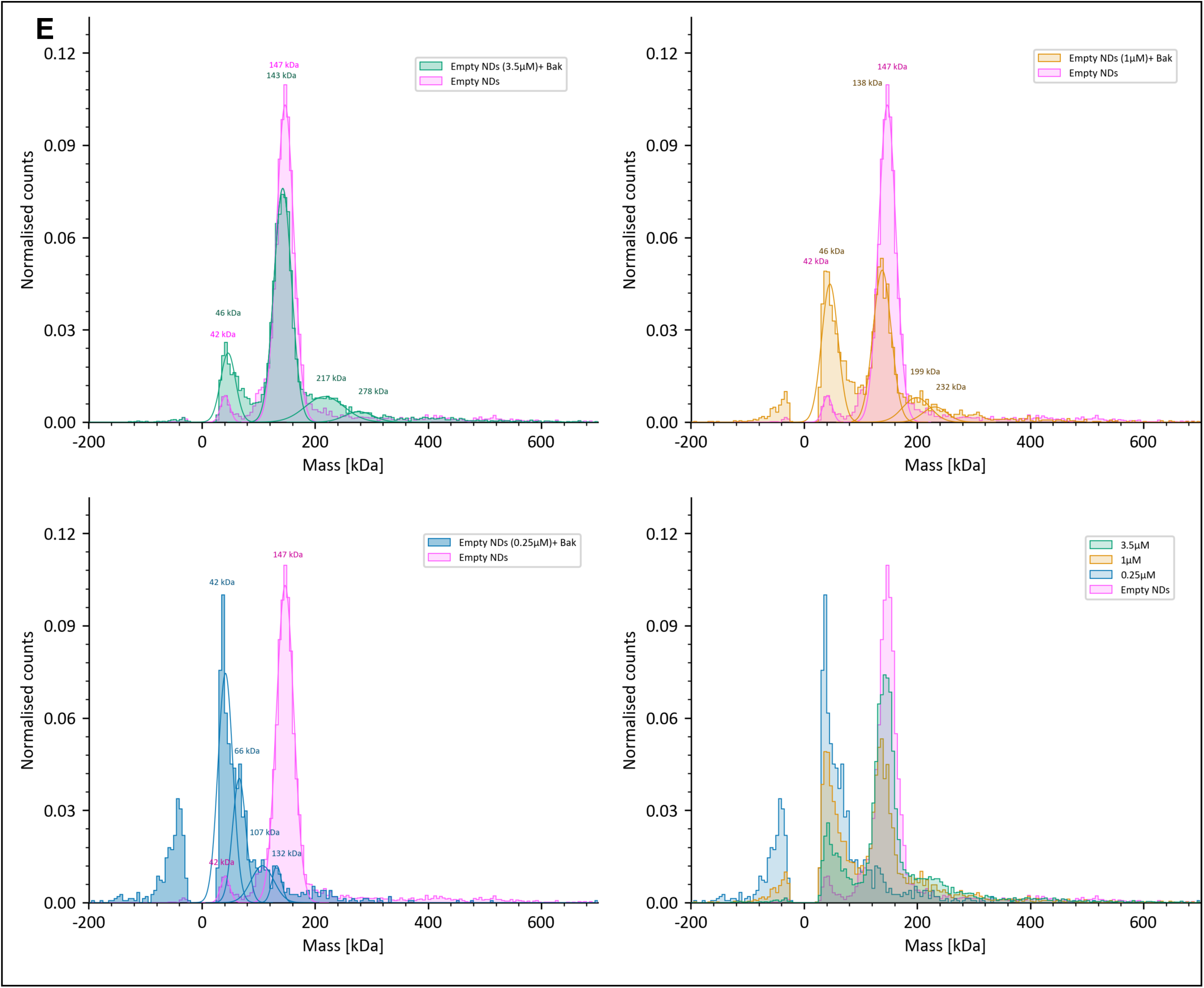
-Supramolecular organization of Bak FL in NDs. A-0.5% of eluates from His-Trap1 untreated or reduced with 2% β-mercaptoethanol (β-ME) were separated by SDS-PAGE followed by immunodetection of Bak. B-Densitometric quantification of n>3 independent experiments. Mean is shown +/-SD, and a statistical Wilcoxon test was used. C-Eluates from His-Trap1were incubated or not with 0.01% Brij 58 prior to cross-linking with 2 mM DSG (disuccinimidylglutarate). 1.5% of E1 was separated on denaturing gradient SDS-PAGE followed by imunodetection of Bak (left) and His7-MSP1E3D1 (right). Arrows indicate homotypic adducts or Bak (cyan) and putative heterotypic Bak-MSP adducts (black). D-Non denaturing PAGE : Bak FL was synthesized in the presence of 3.5 µM NDs and submitted to HisTrap. 2.5% of E1 were separated on gradient native gels, followed by immunodetection of Bak (left) and His7-MSP1E3D1 (right). E-Normalized mass photometry histograms of empty NDs compared to Bak+ND samples obtained after CFPS in the presence of decreasing concentrations of empty NDs (3.5µM, green; 1µM, orange; 0.25µM, blue). Histograms were normalized to the total number of detected events to facilitate comparison between samples. Gaussian fits (solid lines) were used to identify the major mass populations. Empty NDs display a dominant peak centered at approximately 147 kDa. With decreasing concentrations of empty NDs, the relative abundance and distribution of higher-mass species change. The overlay of all normalized distributions (bottom right) highlights the concentration-dependent redistribution of particle masses.

Cross-linking experiments with disuccinimidylglutarate (DSG) performed on E1 fraction further substantiate the close proximity (<7.7 Å) between Bak molecules in NDs. The electrophoretic separation of the DSG-treated sample on gradient SDS-PAGE (Fig. 6C) shows potential homotypic adducts of Bak (bands which appear only on western blots with Bak antibodies but not on western blots with His antibodies), and heterotypic adducts formed between Bak and MSP (bands co-migrating in western blots with Bak antibodies and His antibodies). Homotypic adducts appear at ∼55 kDa (predominant signal), ∼130 kDa and ∼180 kDa, which could respectively correspond to dimers, hexamers and octamers. Putative heterotypic adducts appear at ∼70 kDa and ∼90 kDa. The addition of Brij 58 (a nonionic surfactant) to E1 fractions, prior to DSG treatment, is intended to solubilize NDs and thus prevent the cross-linking of proteins merely juxtaposed in NDs. Only proteins engaged and stabilized in higher order assemblies will survive the solubilisation step. Fig. 6C shows that homotypic adducts of Bak are destabilized and that only Bak dimers remain. We therefore conclude that the stability of Bak higher order assemblies depends on NDs integrity, while Bak dimers are stable and could represent the building unit for oligomers formation.

A complementary technique to evaluate the supramolecular assembly of Bak in NDs was the use of native gels. After CFPS synthesis of Bak FL in the presence of 3.5 µM NDs and HisTrap chromatography, E1 fraction was separated on non denaturing gradient gels (Fig. 6D). The comparison of His signals and Bak signals indicate that under these conditions, NDs are for the most part empty and migrate below 117 kDa (Fig. 6D, right). Bands of much less intensity are visible at higher molecular weight, which co-migrate with bands detected with the Bak antibody (Fig. 6D, left). ND-embedded Bak distribute as discrete assemblies (3 intense bands: one below 171 kDa, one at ∼171 kDa and one above 171 kDa; discrete assemblies of lesser intensity are detectable up to 460 kDa). This indicates that Bak spontaneously forms oligomers up to 8-mers in NDs, but the exact stoichiometry of which will need to be established.

As a way to address the stoichiometry of Bak oligmers in NDs, preliminary biophysical analyses by mass photometry (MP) were carried out on samples where Bak FL was synthesized in the presence of decreasing concentrations of NDs. We reasoned that NDs at 3.5 µM predominantly remained empty, and we therefore also performed CFPS in the presence of 1 µM and 0.25 µM NDs to optimize the ratio of ND-embedded Bak. Fig. 6E shows the MP histogram obtained for each ND concentration, compared to empty NDs for reference. At higher concentrations of NDs (Fig. 6E, top left), the mass distribution is shifted toward species with higher molecular weights, with a lesser relative abundance of low-mass assemblies. The distribution gradually reverses when ND concentration is decreased (Fig. 6E, top right and bottom left), nicely confirming a concentration-dependent redistribution of BAK-containing NDs. Since biochemical analyses have demonstrated the coexistence of monomeric, dimeric, and higher-order BAK oligomers, the broad mass distributions observed by MP most likely reflect heterogeneous occupancies of BAK and oligomeric states within individual NDs, rather than distinct stoichiometric species. These initial results appear encouraging for the use of MP to assess the supramolecular states of BAK and, more broadly, of Bcl-2 proteins in the context of NDs.

## 5. Notes

- ÄKTA chromatography parameters for His-Trap : Auto-zero to set A_280nm_ to 0 mAU / Flow : 1 mL/min / High Pressure limit 0.4 MPa / Fraction size 1 mL.
- ÄKTA chromatography parameters for Superdex 200 10/300 GL increase : Auto-zero to set A_280nm_ to 0 mAU / Flow 0.4 mL/min / High Pressure limit 2.8 MPa / Fraction size 0.5 mL.
- DLS parameters: DLS acquisition time (s)= 5; number of DLS acquisitions = 10; temperature = 25°C; wait = 0; Do = 3;

## 6. Concluding remarks

This work presents a rapid, effective and unified protocol for the production of three different Bcl-2 family proteins by CFPS. This protocol was initially described for Bcl-xL and Bax, and it is now extended to the production of Bak, although for unknown reasons Bak in synthesized in lesser amounts than the other two. These proteins have long been suspected to exhibit remarkable structural plasticity. The present work shows that CFPS offers a valuable technological development to investigate the intrinsic and extrinsic molecular determinants driving the stepwise conformational changes needed to transition from soluble monomeric proteins to membrane-embedded oligomeric proteins. In particular, this work pointed out striking differences between Bax and Bak, where conformational changes of the former are uniquely triggered by the phospholipids of NDs, and where both proteins respond differently to the addition of N-terminal His6-tag. Whether another tag, with a different net charge and different electrostatic properties (for example a Flag-tag), would have the same effect remains an open question.

In spite of the low amount of Bak produced, the method presented herein allows to produce and purify membrane-inserted full-length Bak. We documented that when Bak is synthesized by CFPS in the presence of preformed NDs, it spontaneously inserts in the phospholipid bilayer. The analyses carried out to investigate Bak supramolecular assemblies in NDs show a stable arrangement in dimers, along with an arrangement in oligomers, the stability of which depend on NDs integrity. Furthermore, we demonstrate that the production/purification protocol presented is compatible with sample preparation for high-resolution structural determination. Preliminary data validated the use of mass photometry for the quality control of Bak reconstituted in NDs. Experiments confirmed a clear dependence of Bak assemblies relative to ND concentration; this emphasizes again that the protocol described here does not discard empty NDs, and stresses that Bak to ND ratio will require optimization to increase the likelihood of successful high-resolution structural determination.

Taken together, this work provides the methods to produce membrane-embedded full-length, untagged Bcl-2 protein family. It thus paves the way for long-sought dynamic and structural studies in a membrane environment.

## Acknowledgments

This work was supported by Agence Nationale de la Recherche ANR-22-CE17-0050 to MP. This work was also supported by Société Française d’Hématologie to JK.

